# CRISPR-Cas9 RALA knockout and reconstitution. Detection and role of RALA S194 phosphorylation in RAS-dependent and RAS-independent cancers

**DOI:** 10.1101/2024.12.15.628532

**Authors:** Mayuresh Konde, Siddhi Inchanalkar, Nilesh Deshpande, Tushar Sherkhane, Mishika Virmani, Kajal Singh, Nagaraj Balasubramanian

## Abstract

Downstream of oncogenic RAS, RALA is critical for cancer tumorigenesis, possibly regulated by phosphorylation of its Serine194 residue. We made CRISPR-Cas9 RALA knockout (RALA KO) in three RAS-dependent and two RAS-independent cancer cells. Detection of RALA S194 phosphorylation using the commercial anti-phospho-RALA antibody lacks specificity in all three RAS-dependent cancers. siRNA knockdown of RALA and AURKA inhibition by MLN8237 (V_MLN_) also did not affect pS194RALA detection in these cancers. RALA KO MiaPaCa2 (RAS-dependent) and MCF7 (RAS-independent) cells, stably reconstituted with WT-RALA and S194A-RALA mutants, showed no effect on RALA activation. Tumour growth was, however, restored partly by WT-RALA, but not S194A-RALA mutant. Thus, RALA S194 phosphorylation is needed for tumor formation, not affecting its activation but possibly through its localization.

## INTRODUCTION

RALA regulates cell migration, proliferation, cell spreading, AIG (Anchorage Independent Growth) and tumorigenesis. Its function depends upon its localization and activation in the cell and is mainly regulated by RALGEFs. In recent years, AURKA has emerged as a novel regulator of RALA^1^. AURKA is a well-known mitotic kinase involved in centrosome maturation^2^. In-vitro kinase assay shows it phosphorylates RALA at Serine 194 residue (S194) but not RALB^3^. AURKA-mediated phosphorylation of RALA at S194 regulates its localization at cell membranes^4^. It remains unclear and possibly context-dependent, if AURKA-mediated RALA phosphorylation at S194 can regulate RALA activation. PKA could also mediate RALA phosphorylation at S194^5^.

Studies in non-transformed HEK293 and MDCK cells expressing constitutively active RALA^V23^ and phosphodeficient S194A-RALA^V23^ mutants show this phosphorylation to be vital for RALA-dependent AIG^1,3^. In some pancreatic cancer cells, the knockdown of RALA and reconstitution with WT-RALA and S149A-RALA mutants further support this claim in AIG studies^4^. AURKA inhibitor (MLN8237) treatment of pancreatic and breast cancer cells also affects their AIG^5,6^.

The only known tumor formation study in RALA knockdown pancreatic cancer cells (CFPac-1, HPAC, Capan1) reconstituted with WT-RALA vs S194A-RALA phosphodeficient mutant showed loss of RALA to affect tumor formation that is restored by WT-RALA reconstitution. S194A-RALA mutants, however, had variable effects and were comparable to WT-RALA in CFPac-1, comparable to RALA KD in Capan-1, and better than RALA KD but not equivalent to WT-RALA in HPAC tumors^1^. Targeting AURKA activation using inhibitors like MLN8237 (Alisertib) in pancreatic cancers (PancT4 cell line and Primary cells from tumors) suppresses tumor formation^5^. However, this study showed no change in RALA S194 phosphorylation detected by western blot using anti-phospho-RALA (Merck-Millipore, Cat no-07-2119), suggesting the effects seen might be independent of RALA. These studies were exclusively done in RAS-dependent cancers.

These differences in AURKA-RALA crosstalk observed in non-transformed vs cancer cells could, in part, be mediated by regulatory influencers, like the presence of oncogenic RAS. Oncogenic RAS-mediated regulation of the RAL GTPases is vital for RAS-mediated transformation and is seen to drive AIG^1^. RAS-dependent regulation of RAL is known to be mediated by RAS-dependent RALGEFs^7^. Constitutively active RAS is also seen to promote AURKA expression in cancer cells^8^. RAS is also known to interact with AURKA^9^ and joint expression of both is further seen to promote AIG^10^. This study, hence, evaluates the role RALA S194 phosphorylation has in cancers and what influence the presence of oncogenic RAS could have on its regulation and role. In doing so, it also helps us comment on how the detection of pS194RALA could have implications in interpreting the role of this phosphorylation in cancer cells.

## RESULTS

### CRSPIR-Cas9 mediated stable knockout of RALA

Evaluation of the role of RAL in cancer has been understandably focused on malignancies with RAS mutation, such as lung and pancreatic cancers^11^. Inconsistency in RAL function across cancer types illustrates a need to thoroughly evaluate their contribution and regulation in cancers^11^. CRISPR-Cas9 mediated knockout (KO) poses some advantages to siRNA and shRNA-mediated protein expression targeting, including less off-target effects^12^. Despite some of these advantages, only one previous study in MDA MB231 cells used CRISPR-Cas9 to target RALA and RALB^13^ their role in promoting AIG and tumorigenesis in MDA MB 231 cells.

As a first step to understanding RAS-dependent regulation of RALA S194 phosphorylation and its role in cancers, RAS-independent (MCF7 and SKOV3) and RAS-dependent (T24, UMUC3 and MiaPaCa2) cell lines were chosen based on RALA expression and activation status^1,14,15^. We aimed to use the CRISPR-Cas9 approach to knockout RALA in these cells, and for this, we designed two sgRNAs against exon two and one against exon 4 of RALA using the Chop-chop tool **(Figure 1A)**. *In-silico* checks confirmed their lack of off-target binding and efficiency in making double-strand breaks with Cas9. All sgRNAs were then cloned into the pSpBBCas9-Puro vector. These plasmids were sequenced to confirm the presence of the inserted sgRNAs at the right site **(Supp Figure 1A)**.

**Figure 1:**
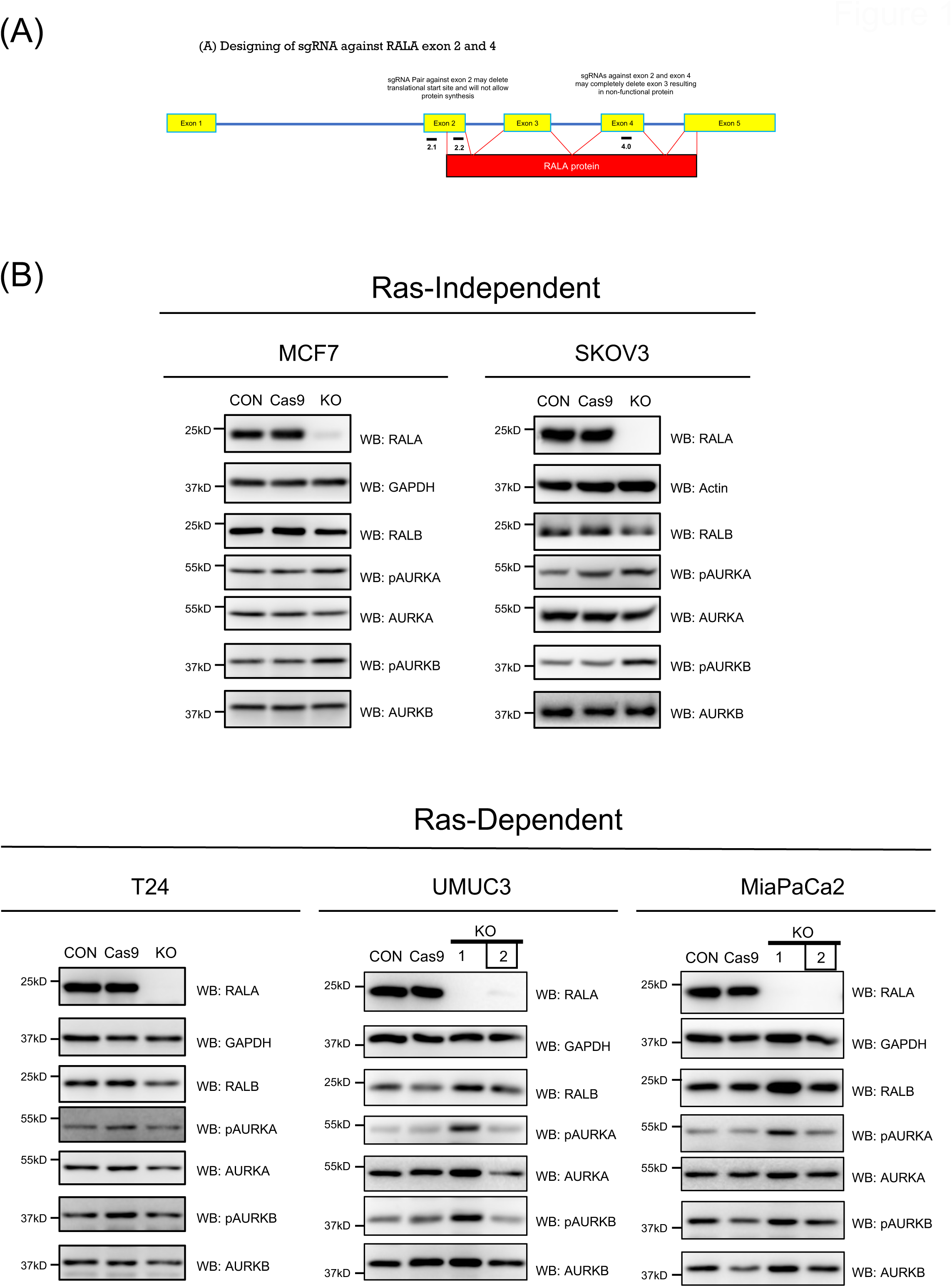
CRISPR-Cas9 mediated knockout of RALA in RAS-independent and RAS-dependent cancer cells. (A) The schematic shows an exon map (yellow boxes) for RALA, marking the location of sgRNAs designed against exon 2 (2.1 and 2.2) and exon 4 (4.0) to target RALA. The red box represents the functional RALA protein coded by these exons. (B) Western blot detection (WB) of RALA, RALB, AURKA, AURKB in untreated control (CON), Cas9 control (Cas9), RALA knockout (KO) clones of RAS-independent (MCF7 and SKOV3) (single clone each) and RAS-dependent (T24, UMUC3 and MiaPaCa2) (two clones for some) cells. Clone 2 selected based on these studies is marked by a BOX. This was accompanied by the Western blot detection (WB) of active AURKA (pAURKA – pThr288 AURKA) and active AURKB (pAURKB – pThr232 AURKB).

Cancer cells were transfected with all three sgRNAs and the pSpBBCas9-GFP vector (as a marker for transfection), single-cells were sorted, clones expanded and tested for RALA expression. Preliminary screening for RALA vs RALB expression allowed us to select clone(s) with the best targeting of RALA without affecting RALB **(Figure 1B)**. One clone, each for MCF7, SKOV3 and T24 and two clones for UMUC3 and MiaPaCa2, were then probed for the expression of RALA, RALB, and regulatory players that could be affected by this targeting **(Figure 1B)**. Together, this confirmed that the loss of RALA in MCF7, SKOV3, T24, UMUC3 (clone 2), and MiaPaCa2 (clone 2) without affecting the expression of RALB, AURKA, AURKB **(Figure 1B)**, AKT, ERK and AMPK **(Supp Figure 1B)**. No detectable change in the activation of AURKA, AURKB **(Figure 1B)**, AKT, ERK and AMPK **(Supp Figure 1B)** was seen, making them suitable for further evaluation.

PCR using genotyping primers to detect a physical break in exon 2 and/or exon 4 of RALA (reflected as a loss or shift in the size of the PCR amplicon) confirmed the KO status of the selected clones for MCF7, T24, UMUC3 (clone 2) and MiaPaCa2 (clone 2) **(Supp. Figure 1C)**. MCF7, T24, UMUC3 and MiaPaCa2 showed a cut in exon 2, while MCF7 and MiaPaCa2 also had a cut in exon 4. The lack of a detectable loss or shift in amplicon for UMUC3 (clone 1) and MiaPaCa2 (clone 1) ruled them out for further use **(Supp. Figure 1C)**. Despite multiple attempts with multiple primers for genotyping PCR in SKOV3 cells, no amplicon was detected. This inexplicable outcome also ruled out using SKOV3 in further studies **(Supp Figures 1C)**. This now gives us a confirmed RALA KO clone for MCF7 (RAS-independent) and T24, UMUC3, and MiaPaCa2 (RAS-dependent) cell lines to take forward in these studies. With issues in genotyping (as seen for SKOV3) and non-specific targeting of genes^16^, validating CRISPR-Cas9 knockouts becomes particularly important with only one previous study in MDA MB231 cells using CRISPR-Cas9 to target RALA^13^, the clones selected and verified to constitute a strong basis for further evaluation of the role of RALA and its mutants.

### Detection of RALA S194 phosphorylation in RAS-independent and RAS-dependent cancer cell lines

Using the RALA knockout clones, we hoped to establish the specificity of the extensively used anti-phospho-RALA Antibody (Serine 194) (Merck-Millipore, Cat no-07-2119) raised against KLH-conjugated linear peptide corresponding to human RALA phosphorylated at Serine 194. This antibody is used across different studies^5^ and has been reported to occasionally detect pS194 RALA in cancer cells despite the loss of RALA ^5^. This had raised questions about its specificity.

Using cell lysates from confirmed RALA KO clones of MCF7, T24, UMUC3 and MiaPaCa2 (evaluated in Figure 1) were probed with the anti-phospho-RALA Antibody and found its detection of pS194RALA variable. No detectable band for pS194 RALA was seen in RALA KO RAS-independent MCF7 cells **(Figure 2A)**. In SKOV3 cells, despite genotyping issues, a near-complete loss of RALA was detected, which also showed no detectable band for pS194 RALA **(Figure 2A)**. However, in RAS-dependent T24, UMUC3 and MiaPaCa2 cells, despite the visible absence of RALA in the KO clones, a distinct band at the expected size was consistently detected by the pS194 RALA antibody **(Figure 2A)**. This was comparable in molecular weight and intensity to the pS194RALA band detected in their control cells. To further confirm this observation, we used RALA-specific siRNA^14^ to knock down RALA in MiaPaCa2 cells. Despite a ∼80% drop in RALA protein levels, a distinct band at the expected size was detected by the phospho-RALA antibody, comparable to control **(Figure 2B)**.

**Figure 2:**
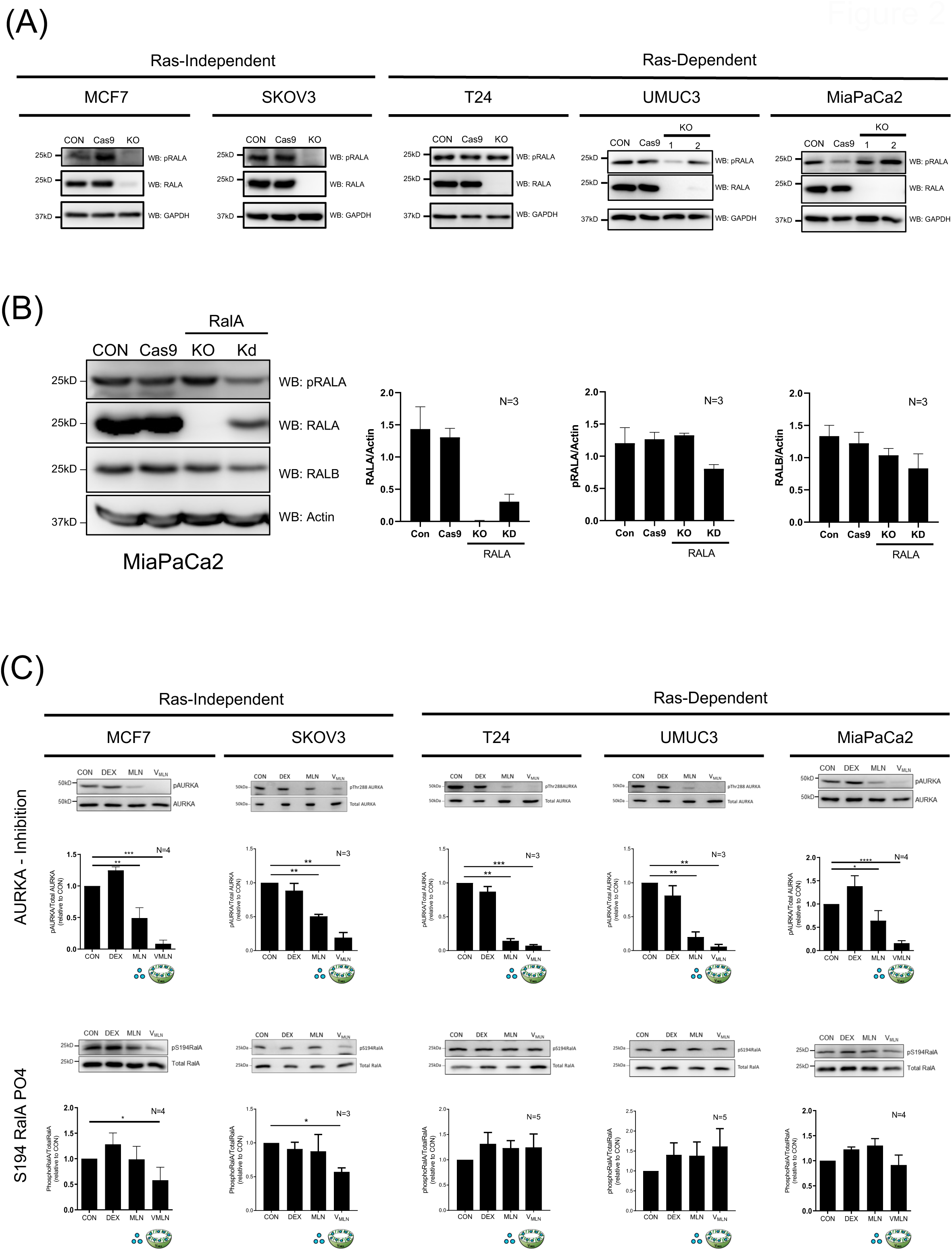
pS194 RALA detection in RAS-independent and RAS-dependent cancer cells. **(A)** Western blot detection of pRALA (RALA S194 phosphorylation) in cell lysate from untreated control (CON), Cas9 control (Cas9) and RALA KO cells using anti-phospho-RALA Antibody (Ser194) (Merck-Millipore, Cat no-07-2119) in RAS-independent **(MCF7 and SKOV3)** and RAS-dependent **(T24, UMUC3 and MiaPaCa2)** cancer cells. **(B)** Western blot detection of pRALA (RALA S194 phosphorylation) in the cell lysate of control (CON), Cas9 control (Cas9), siRNA mediated RALA KD and CRISPR-Cas9 RALA KO MiaPaCa2 cells. Blots were also probed for RALA, RALB and actin. Blots were quantitated, and the ratio of band intensities for RALA, pRALA and RALB normalized to actin as mean ± SE from three independent experiments. Statistical analysis was done using the Mann-Whitney test, compared to control p values, if significant, are represented in the graph (* p < 0.05, ** p<0.01, *** p<0.001). **(C)** Western blot detection (upper panel) and quantitation (lower panel) of phosphorylation of Threonine 288 residue (pThr288) AURKA, total AURKA, phosphorylation of RALA Serine 194 residue (pRALA) and total RALA, from lysates of MCF7, SKOV3, T24, UMUC3 and MIAPaCa2 treated for 48hours with DMSO (CON) and empty nano-vesicle scaffold (DEX), 0.02μM Free MLN (MLN) and 0.02μM encapsulated MLN (V_MLN_). The ratio of pAURKA/AURKA and pRALA/TotalRALA in each cell line was normalized to their respective controls (CON) (equated to 1) and were represented in the graph as mean ± SE from at least three independent experiments. Statistical analysis was done using one sample T-test, compared to control p values, if significant, are represented in the graph (* p < 0.05, ** p<0.01, *** p<0.001).

As an additional approach to confirm this specificity issue, we tested the effect inhibition of AURKA (known to phosphorylate RALA at S194) has on its detection. Previous studies from the lab have developed a dextran nano-vesicle encapsulated MLN8237 (V_MLN_), which inhibits AURKA without affecting AURKB^6^. V_MLN_ uptake was confirmed in MCF7, SKOV3, T24, UMUC3 and MiaPaCa2 cells using Rhodamine (RhB) encapsulated nano-vesicles **(Supp. Figure 2A)**. V_MLN_-mediated inhibition of AURKA (pThr288) was better than free MLN8237 in these cell lines **(Figure 2C)**. Interestingly, AURKA inhibition causes a significant decrease in detectable pS194RALA in MCF7 and SKOV3 cells but not in T24, UMUC3 and MiaPaCa2 cells relative to control cells **(Figure 2C)**. Together, these observations indeed question the specificity of phospho-RALA antibodies in RAS-dependent (T24, UMUC3 and MiaPaCa2) cells. It would, however, be premature to assume that this difference in the detection of pS194RalA depends on the presence of oncogenic RAS, and further evaluation is needed to confirm this. This finding does, however, suggest western blot detection of pS194RalA with this commonly used antibody could be fraught with issues, which could impact experimental outcomes, affecting how we and other studies interpret changes (or lack thereof) in pS194RALA levels in cells.

Earlier studies from the lab showed RALA activation to be regulated by cell-matrix adhesion^14,17^. Serum-deprived mouse fibroblast, when held in suspension for 90 minutes, showed a distinct drop in RALA activation that recovers on re-adhesion **(Figure 3A)**. Interestingly, AURKA activation (pThr288 AURKA) in these cells increases the loss of adhesion and is restored on re-adhesion **(Figure 3B)**. This suggests that AURKA and RALA activation are inversely correlated here. Knowing the anti-phospho-RALA Antibody to be specific to human RALA, we expressed human RALA in mouse fibroblast cells. We detected a distinct increase in S194 RALA phosphorylation on the loss of adhesion, which is restored on re-adhesion **(Figure 3C)**. This suggests that the adhesion-dependent regulation of AURKA controls RALA phosphorylation (pS194 RALA), which negatively regulates RALA activation in response to loss of adhesion **(Figure 3)**. Earlier studies have shown RALA phosphorylation to be a positive regulator of RALA activation in MDCK (Wu et al., 2005) and HEK-TtH cells^1^. They suggest that AURKA-RALA crosstalk and its effect on RALA activation could be cell-type and context-dependent. The lack of antibody specificity in detecting pS194 RALA in some cancer cells can further complicate these interpretations. The most effective way to overcome these limitations and test the role of RALA S194 phosphorylation in RALA function would be to evaluate RalA KO cells expressing RALA phosphomutant (S194A and S194D).

**Figure 3:**
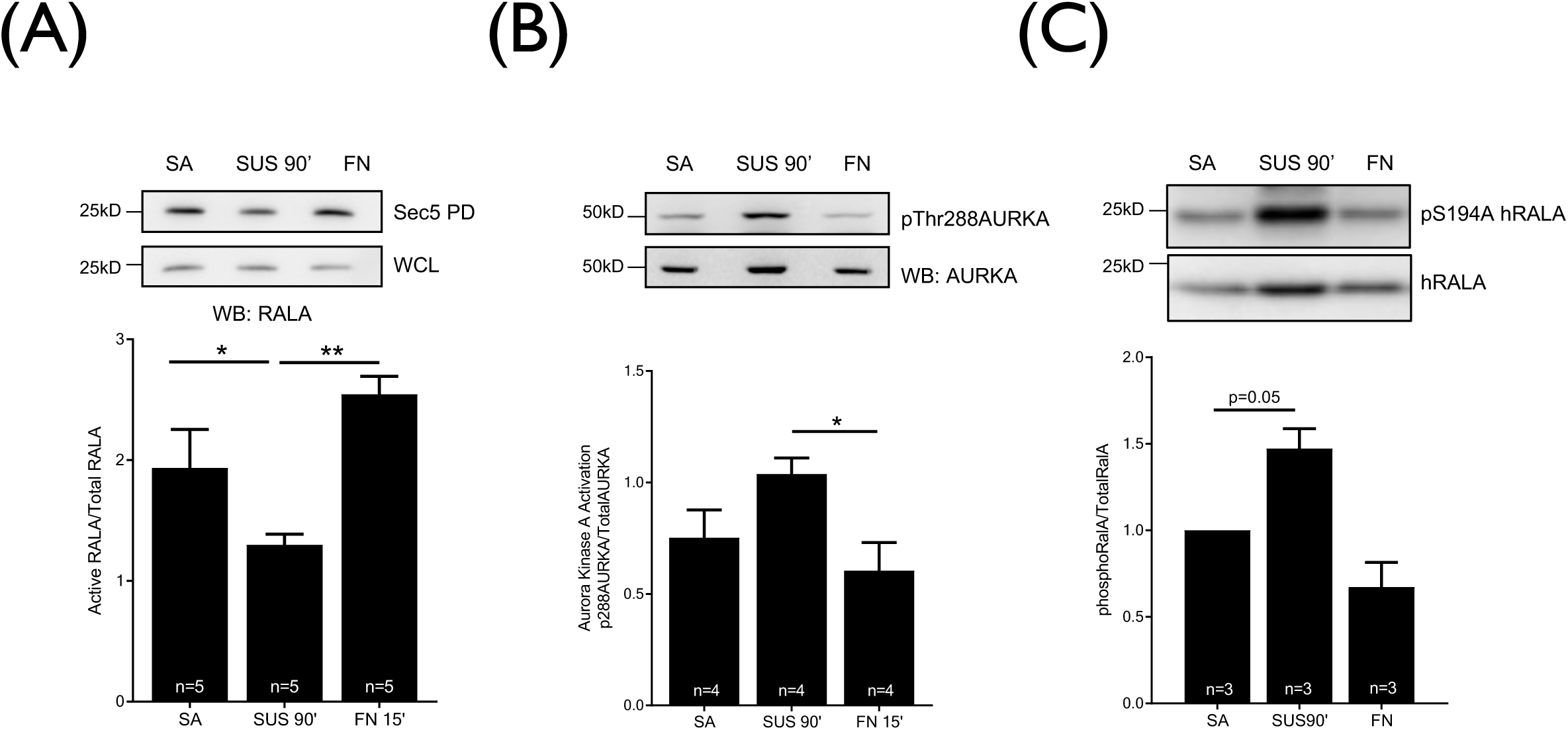
Regulation of RALA and AURKA in early adhesion in WTMEFs. Western blot detection (upper panel) and quantitation (lower panel) of **(A)** Active RALA pulled down by GST-Sec5 (Sec 5), and the total RALA in whole cell lysate (WCL) from serum-starved WT-MEFs that are stable adherent (SA), suspended for 90 min (SUS 90’) and re-adherent on fibronectin for 15mins (FN 15’) probed for RALA (WB : RALA). The graph represents the percentage of active RALA calculated as described in methods from four independent experiments. **(B)** These lysates were probed to detect the phosphorylation of Threonine 288 residues of AURKA (pThr288AURKA) and total AURKA. **(C)** Cell lysates made from human RALA expressing serum starved WT-MEFs stable adherent (SA), suspended for 90 min (SUS 90’) and re-adherent on fibronectin for 15mins (FN 15’) were probed for phosphorylation on Serine 194 residues of human RALA (pSer194hRALA) and total expressed human RALA from four independent experiments. The graph represents the mean ± SE. Statistical analysis of data normalized to CON was done using the paired t-test otherwise was done using Mann-Whitney test, and p values, if significant, are represented in the graph (* p < 0.05).

### Role of RALA and its phosphorylation in RALA activation

Verified RALA KO clones for MCF7 and MiaPaCa2 (clone 2) were used to make stable reconstitutions with WT-RALA, S194A-RALA (phosphodeficient) and S194D-RALA (phosphomimetic). While the S194A-RALA mutant effectively blocks phosphorylation in previous studies^3,5^, the S194D mutant, when used, has been reported to behave like the S194A mutant^1^. These RALA mutants were made in the lab and confirmed by sequencing **(Supp Figure 3A)** and PCR amplification **(Supp Figure 3B)**. In MCF7 KO cells, WT-RALA expressed is detected by the phospho-RALA antibody, while the RALA mutants (S194A/S194D) do not confirm the mutation of the S194 site **(Supp Figure 3C)**.

Retroviral stable expression of WT-RALA and RALA mutants (S194A/S194D) in RALA KO MCF7 and MiaPaCa2 cell lines were comparable when detected with total RALA antibody **(Figure 4A and 4B)**. Their expression was, however, less than endogenous RALA in control cells (CON) **(Figure 4A and 4B)**. GST-Sec5 pulldown of active RALA from reconstituted MCF7 and MiaPaCa2 cells showed no significant difference in the activation status of WT-RALA, S194A-RALA and S194D-RALA mutants **(Figure 4C and 4D)**. This suggests that RALA S194 phosphorylation does not affect RALA activation in RAS-independent (MCF7) or RAS-dependent (MiaPaCa2) cancer cell lines. A small non-significant increase in RALB activation was seen upon RALA KO in both MCF7 and MiaPaCa2 **(Figure 4E and 4F)**, which was unaffected by RALA mutant reconstitution **(Figure 4E and 4F)**.

**Figure 4:**
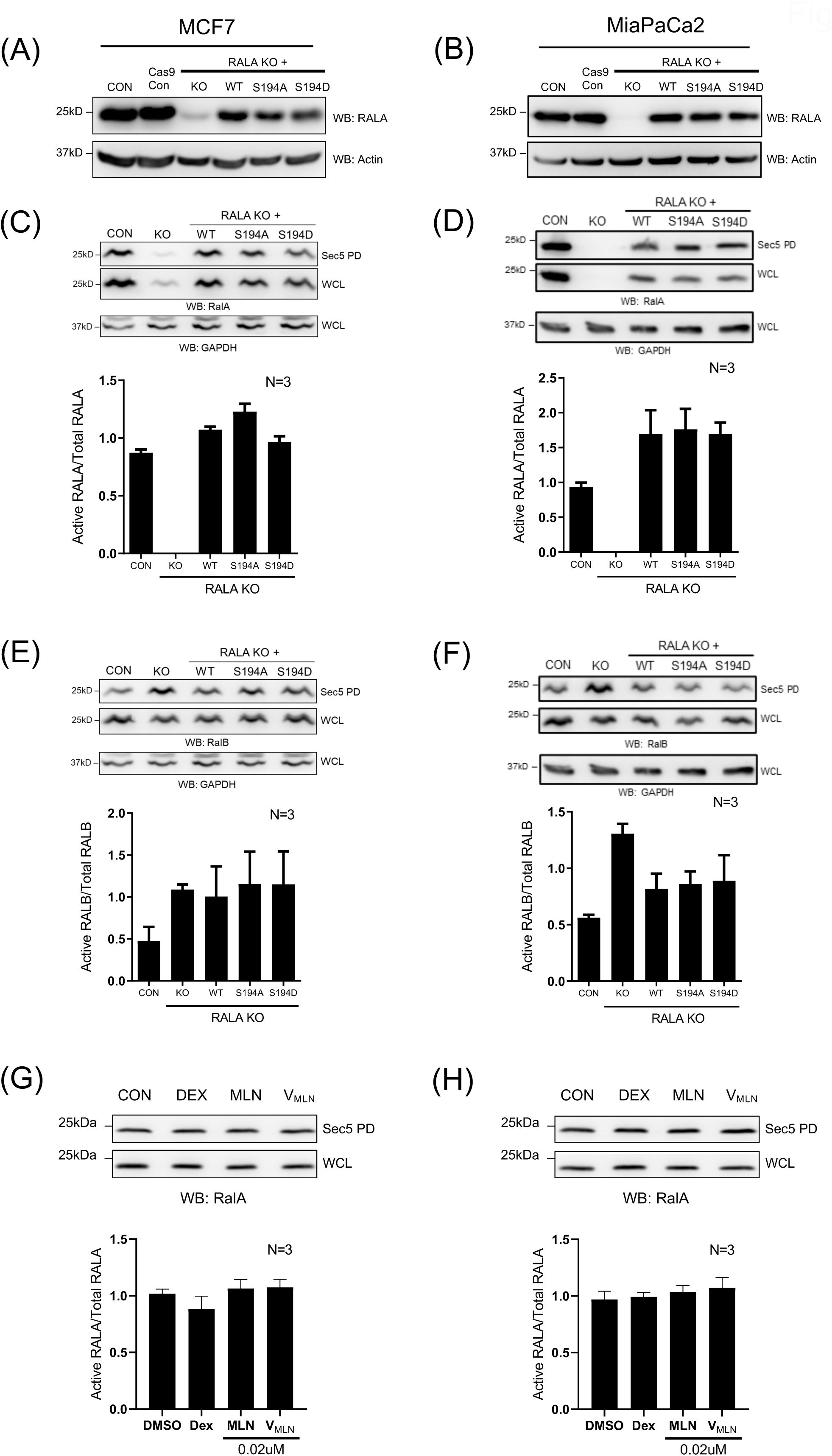
Role of RALA and its S194 phosphorylation in regulating RAL activation in MCF7 and MiaPaCa2 cells. **(A,B)** Western blot detection of RALA (WB : RALA) and actin (WB: Actin) in cell lysates of MCF7 and MiaPaCa2 control (CON), RALA KO (KO) and RALA KO cells reconstituted with RALA mutants (WT/S194A/S194D). **(C,D,E,F)** Western blot detection (upper panel) and quantitation (lower panel) of Active RALA, active RALB, total RALA and total RALB from lysates of Control (CON), Cas9 control (Cas9), RALA KO (KO), WT-RALA (WT), S194A-RALA (S194A) and S194D-RALA (S194D) mutant reconstituted MCF7 and MiaPaCa2 cells. **(G,H)** Western blot detection (upper panel) and quantitation (lower panel) of Active RALA and total RALA from lysates of MCF7 and MiaPaCa2 cells treated with DMSO (CON), empty nano-vesicle scaffold (DEX), 0.02μM Free MLN and 0.02μM V_MLN_. The graphs represent the ratio of active to total protein (RALA and RALB) as mean ± SE from at least three independent experiments. Statistical analysis was done using the Mann-Whitney test, and p values, if significant, are represented in the graph (* p < 0.05, ** p<0.01, *** p<0.001).

As an additional means of confirming if RALA phosphorylation does not affect RALA activation in these cancers, we asked if V_MLN_-mediated inhibition of AURKA (seen to affect RALA phosphorylation in MCF7 vs MiaPaCa2 cells differentially) **(Figure 2C)** regulates RALA activation differently **(Supp Figure 3D)**. When compared to respective controls (CON, DEX), V_MLN_ and free MLN8237 (MLN) treatment did not affect RALA activation in MCF7 or MiaPaCa2 cells **(Figure 4G and 4H)**. This agrees with the RALA mutant data in both cell types. There is the possibility that the lack of a visible change in pS194RalA levels in MiaPaCa2 on V_MLN_ treatment may reflect the lack of specificity in the antibody detection in these cells.

In earlier studies, the regulation of RALA activation by AURKA is variable, with WT-AURKA expression increasing RALA S194 phosphorylation and activation^1^, with kinase-active AURKAThr288 only further increasing RALA S194 phosphorylation without affecting RALA activation^1^. In HEK-TER and NIH3T3 cells, the RALA S194A mutant showed a drop in RALA activity^1,18^. With no known studies evaluating RALA phosphomutants in cancer cells, this study data, for the first time, indicates that RALA S194 phosphorylation does not affect RALA activation in cancer cells.

### Role of RALA and its phosphorylation in tumor formation

The best evaluation of RALA function in cancers stems from its ability to support tumor formation. MCF7 and MiaPaCa2 cells on RALA KO form significantly smaller tumors than control cells **(Figure 5A, 5B)**. This suggests the tumorigenesis of both these cancer cells to be RALA-dependent. Evaluating the tumor-forming ability of WT-RALA expressing MCF7 and MiaPaCa2 cells showed a significant increase in tumor size, confirming the role of RALA in their tumorigenesis **(Figure 5A, 5B)**. This rescue being partial could, in part, reflect the reduced WT-RALA levels in reconstituted KO cells (MCF7 and MiaPaCa2) when compared to endogenous RALA (CON and Cas9-CON) **(Figure 5A, 5B, 5C, 5D)**.

**Figure 5:**
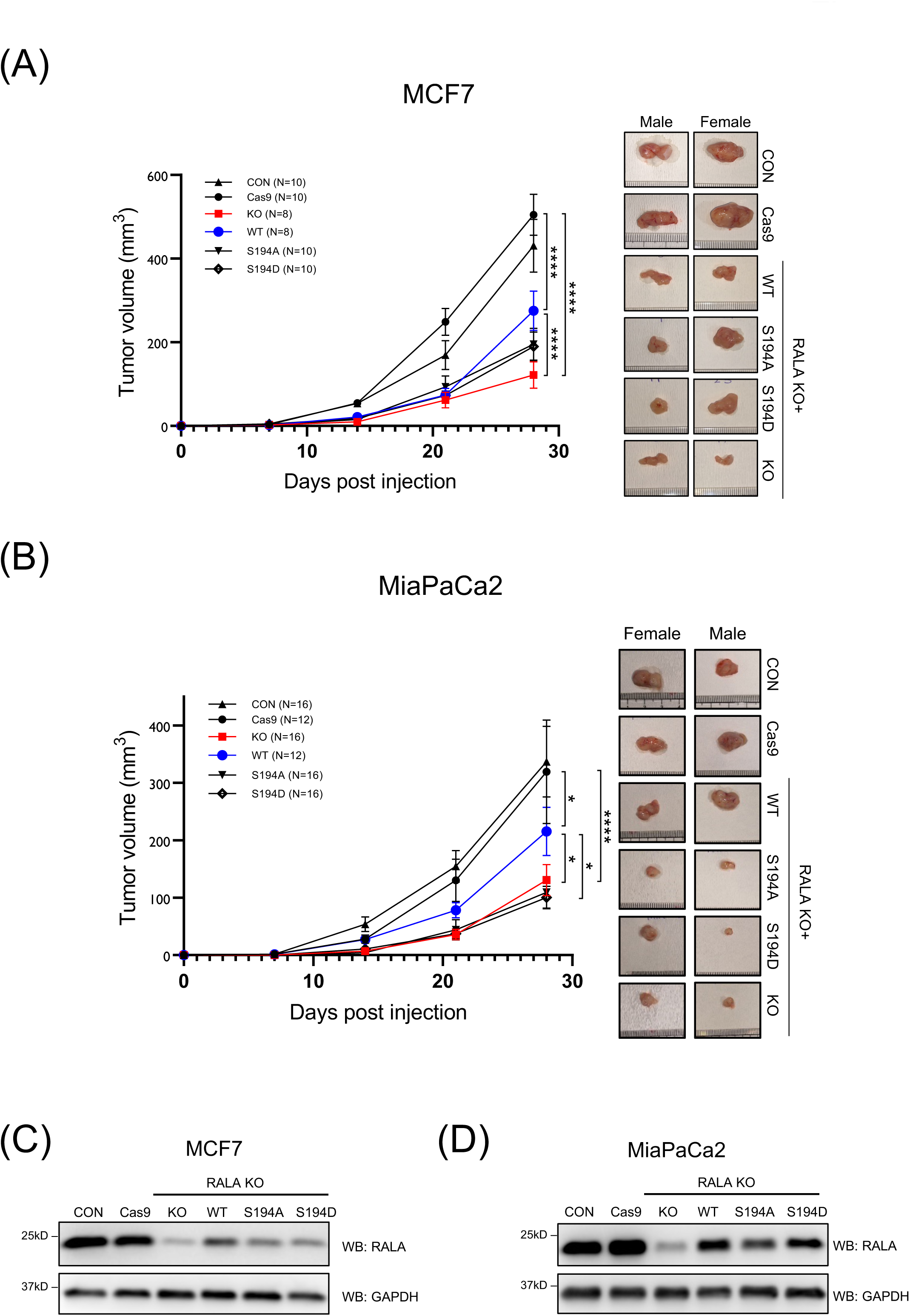
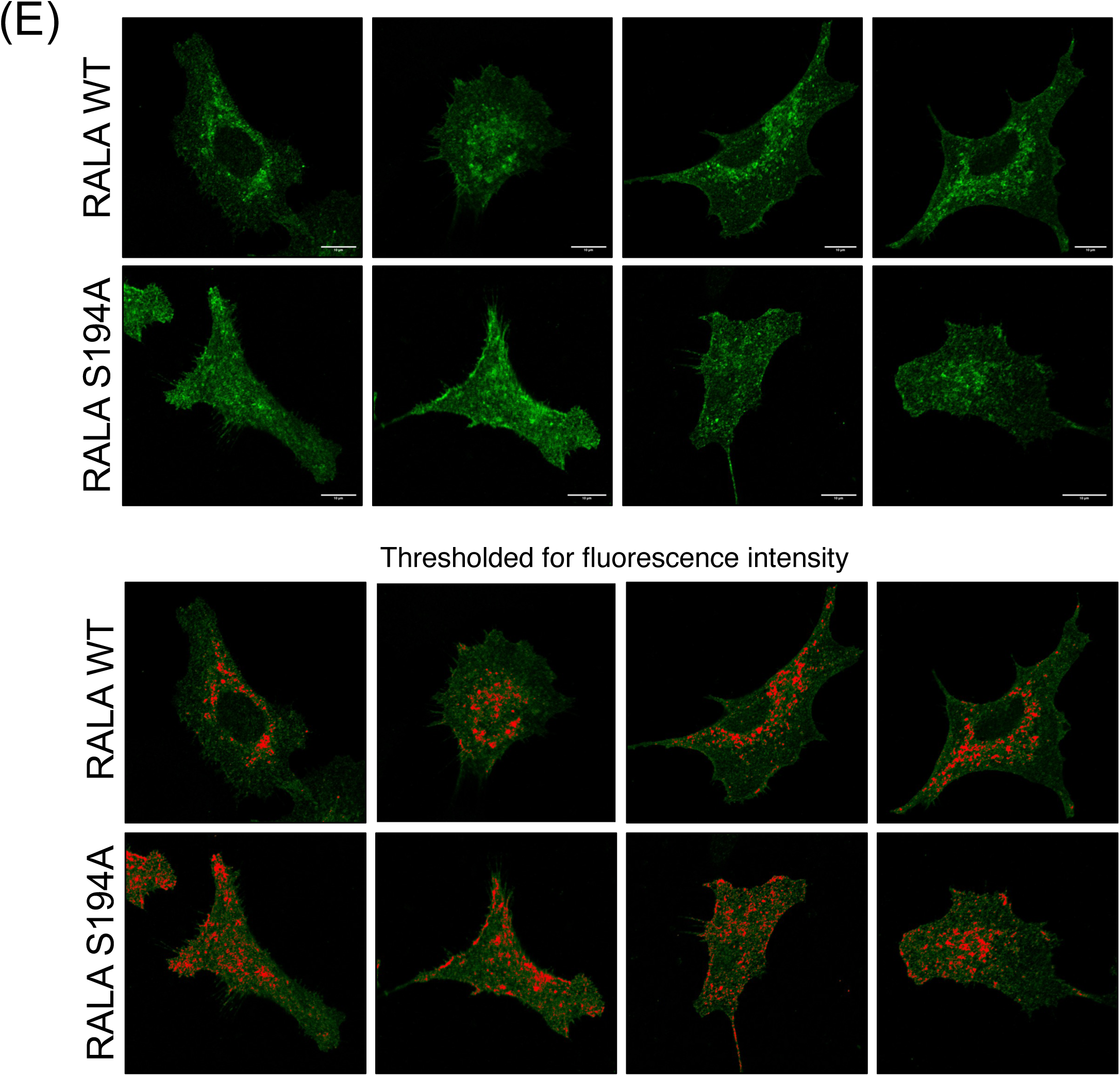
Role of RALA and its S194 phosphorylation in regulating tumor formation and RALA localization in MCF7 and MiaPaCa2 cells. Graph showing mean tumor volume (mm^3^) ± SE over 28 days for tumors made by MCF7 (A) and MiaPaCa2 (B) control cells (CON) (▴), Cas9 treated cells (Cas9) (●), RALA KO cells (KO) (▪) or RALA KO cells expressing WT-RALA (WT) (●) or S194A-RALA (S194A) (▾) or S194D-RALA (S194D) (♢) mutant cells. Statistical analysis was done using Two-Way ANOVA comparing all tumors and p values if significant are represented in the graph (* p < 0.05, **** p < 0.0001). (C,D) Western blot detection of RALA and Actin from harvested tumor lysates (protein equivalent) of control cells (CON), Cas9 control (Cas9), RALA KO (KO) and RALA mutants (WT, S194A and S194D) in MCF7 and MiaPaCa2 cells. (E) Immunofluorescence detection of RALA in MCF7 RALA KO cells reconstituted with WT-RALA (top panel) and S194-RALA mutant (bottom panel). Fluorescence images were comparably thresholded, and intensities above the set threshold were marked in red to highlight their localization.

Tumors formed by MCF7 and MiaPaCa2 S194A-RALA mutant reconstituted cells were comparable to their RALA KO tumors in size **(Figure 5A, 5B)**. They show that RALA S194 phosphorylation is indeed required for RALA-dependent tumor formation. In RAS-dependent MiaPaCa2 cells, the S194A-RALA mutant cells behaved similarly to the RALA KO cells in the size of their tumors **(Figure 5B)**. In RAS-independent MCF7 cells, the S194A-RALA mutant cells behaved slightly differently. Their tumor sizes were marginally (not significantly) bigger than the RALA KO tumors and marginally (not significantly) smaller than the WT-RALA reconstituted tumors **(Figure 5A)**. This suggests that while tumor formation in MiaPaCa2 and MCF7 are dependent on RALA phosphorylation, their relative contribution could be variable between tumors, as reported earlier in pancreatic cancer cells^1^.

In both MCF7 and MiaPaCa2 cells, tumors formed by S194D-RALA mutant reconstituted cells were comparable in their size to the S194A-RALA mutants **(Figure 5A, 5B)**. This could mean the S194D-RALA mutant may not effectively mimic RALA phosphorylation in these cells. An earlier study with the S194D mutant reported that it behaves similarly^1^. This limits its use in evaluating the role of this phosphorylation in RALA function.

The localization of RALA and its regulation by RALA S194 phosphorylation could, in the absence of any effect on RALA activation, be vital for regulating tumor growth. In MCF7 cells, the WT-RALA showed a prominent perinuclear and occasional membrane ruffle localization. S194A-RALA mutant lacked this spatial clarity in cells, being more uniformly distributed throughout the cell. When comparably thresholded for intensities, fluorescence images clearly captured this spatial difference **(Figure 5E)**. This localization difference in MiaPaCa2 cells was difficult to evaluate as cells spread distinctly less. Earlier studies in RALA knockdown HEKTtH cells reconstituted with WT-RALA vs S194-RALA mutant suggest a change in their localization to internal membranes, as seen in fractionation studies. S194 phosphorylation is seen to influence the interaction of RALA effector RALBP1^1^. Aurora A phosphorylated RALA, is known to relocalize to the mitochondria, where it is seen to concentrate RALBP1 and DRP1to regulate mitochondrial fission^19^. The implication this could have in tumors remains unclear. Could a phosphorylation-mediated change in RALA localization affect its interaction with downstream effectors^1^ to impact its role in tumor formation is a question of much interest.

Together, these studies implicate S194 phosphorylation as vital for the role RALA has in tumor formation in MCF7 and MiaPaCa2 cells, independent of its activation and possibly regulated by its localization. Since S194 phosphorylation is vital for RALA-dependent tumor formation, targeting this site can have therapeutic advantages. Could AURKA targeting be an effective way to block RALA-dependent tumor growth is, hence a question worth addressing.

## ACKNOWLEDGMENTS

We thank Prof. Manickam Jayakannan, Department of Chemistry, IISER Pune for valuable discussion and nano-vesicle making. This work was supported by ICMR grant 35/03/2019-NANO/BMS. Mayuresh and Nilesh were supported by UGC fellowship, Siddhi was supported by DBT fellowship, Kajal was supported by CSIR fellowship. We also thank IISER Pune Microscopy facility, Flow cytometry facility and National Facility for Gene Function in Health and Disease (NFGFHD) facility for their support.

## MATERIALS AND METHODS

### Cell culture and reagents

MCF7, SKOV3, T24, UMUC3, MiaPaCa2 and HEK293T cells were obtained from ATCC or ECACC. SKOV3, UMUC3 and T24 cells were cultured in high glucose DMEM medium with 5% fetal bovine serum (FBS) and MCF7 and MiaPaCa2 cells were cultured in high glucose DMEM medium with 5% fetal bovine serum (FBS), penicillin, and streptomycin (Invitrogen).

Human plasma fibronectin (Cat#F2006), nocodazole (Cat # M1404), DMSO (Cat # D2438), DAPI (Cat# D9542), sodium orthovanadate (Cat# S6508), sparfloxacin (Cat# 56968) and thiazolyl blue tetrazolium bromide (MTT, Cat# M2128) were purchased from Sigma, and Phalloidin-Alexa-488 (Cat# A12379), Phalloidin-Alexa-633 (Cat # A22284) was from Molecular Probes (Invitrogen). Fluoromount-G was used to mount cells for imaging and was obtained from Southern Biotech (Cat# 0100-01). Glutathione sepharose beads used for the Ral activity assay were from GE (Cat# 17075601). Crystal violet was used for staining colonies in the AIG assay from Amresco (Cat# 0528). Alisertib (MLN8237) was purchased from Selleckchem (Cat# S1133). BQU57 (Ral inhibitor Cat# SML-1268), and AZD1152 (AURKB inhibitor Cat# SML0268) were purchased from Sigma. The BCA protein estimation kit (Cat# 23227) was purchased from Thermo Scientific. Nonidet P40 Substitute (Cat# 68387-90-6) and RNAase-A (Cat# 9001-99-4) were purchased from USB corporation. Trizol (Cat# 15596018) was purchased from Ambion. Immobilon western blot substrate (Cat# WBKLS0500) was purchased from Millipore.

Antibodies used for western blotting include anti-phospho-aurora A (Thr288)/aurora B (Thr232)/ aurora C (Thr198) (Cat #2914), anti-AURKB (Cat #3094), anti-phospho AKTSer473 (Cat# 9271) (1:2000 dilution), anti-phospho-FAK-Tyr397 (Cat# 3283), anti-phosphop44/ p42 ERK1/2 (Thr202/Tyr204) (Cat# 4370) (1:2000 dilution), anti-p44/p42 ERK1/2 (Cat# 4695) (1:2000 dilution), anti-FAK (Cat#3285), anti-AKT (Cat# 4691) antibodies were purchased from Cell Signaling Technologies and used at 1:1000 dilution unless mentioned otherwise. Anti-AURKA (Cat #610939) and anti-RALA (Cat# 610221, clone 8) were purchased from BD Transduction Laboratories and used at 1:1000 dilution. Anti-Phospho-RALA (Ser194) (Cat# 07-2119) was purchased from Millipore, and Anti-RALB was from R&D Laboratories (Cat# AF3204), both used at 1:1000 dilution. Anti-beta actin (Cat# Ab3280) antibody was purchased from Abcam and used at 1:2000 dilution. Secondary antibodies conjugated with HRP were purchased from Jackson Immunoresearch and were used at a dilution of 1:10000.

### Making of Nanovesicle Encapsulated MLN8237 (V_MLN_) with Rhodamine B

Amphiphilic dextran (DEX-PDP) was synthesized as reported earlier (Pramod et al. 2012) and used to encapsulate MLN8237 in the polysaccharide (dextran) vesicles as reported earlier by (Inchanalkar et al. 2018). The drug loading content (DLC) and drug loading efficiencies (DLE) were determined by dissolving a known amount of a lyophilized drug-loaded sample in methanol and estimating its drug content by absorption spectroscopy. The molar extinction coefficient of MLN8237 was determined as 76500 L mol−1 cm−1 in methanol. The DLC and DLE were determined as 0.40% and 56%, respectively.

### Cellular Uptake of V_MLN_ by Confocal Microscopy

MLN8237 and Rh-B were encapsulated together in dextran vesicles as described by (Inchanalkar et al. 2018) and their DLCs and DLEs were determined to be 0.42% and 58% for MLN8237 and 1.4% and 64% for Rh-B, respectively. Incubating cancer cells with V_MLN,_ their cellular uptake was evaluated in detail earlier (Inchanalkar et al. 2018).

### MLN8237 and V_MLN_ mediated AURKA inhibition

Cells (0.5 ×10^5^ cells) were seeded in 6 well plates for 24 hours, and treated with 0.02 μM MLN8237 as a free drug or in the nanovesicle (V_MLN_) for 48hours. In all inhibition experiments, DMSO (CON) and empty dextran scaffold (DEX) were used as were solvent controls for free MLN8237 and encapsulated MLN8237 (V_MLN_), respectively. At the end of 48 hours, cells were lysed in 150ul of Laemmli buffer, resolved by SDS PAGE and western blotting was performed to check pAURKA, AURKA, pS194RALA and RALA.

### RAL activity assay

Control, RALA KO and RALA mutant reconstituted cells (2.5 × 10^5^) were seeded in a 10cm dish for 72 hours with or without nanovesicle-encapsulated inhibitor (V_MLN_). As reported earlier, active RAL was pulldown with the GST-Sec5 Ral binding domain (RBD) (Inchanalkar et al. 2018). To determine the percentage of active RALA and RALB, the following calculation was used:

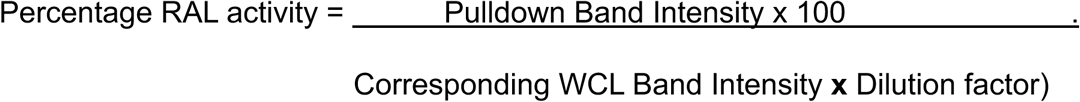

The dilution factor was calculated as the ratio of the amount of total cell lysate used for the pulldown (400 μl) and the amount of this lysate resolved by SDS PAGE as the whole cell lysate (WCL) (24 μl WCL + 6 μL 5X Lamelli buffer). The dilution factor was hence 400 ÷ 24 = 16.66. This was constant in all experiments.

### Anchorage-Independent Growth (AIG) Assay

The effect of AURKA inhibition by a nanovesicle-encapsulated inhibitor (V_MLN_) on AIG was tested as described earlier (Inchanalkar et al. 2018). Cells were treated with V_MLN_ at 0.02uM concentration for 10-14 days with fresh V_MLN_ added every 3^rd^ day.

### Making of CRISPR-Cas9 KO in cancer cell lines

We designed 3 sgRNAs against RALA, two were targeted against exon 2, flanking the TSS (Translation start site) and one against exon 4. CHOP-CHOP and CRISPR KO online tools were used to design sgRNAs **(as described in Table 1)** with high efficiency and low off-target binding. Low off-target binding was confirmed with the CCTOP online tool. sgRNAs were cloned into the pSpBBCas9-Puro vector using a restriction-based cloning method, as mentioned by Ran et al. 2013.

**Table 1.**
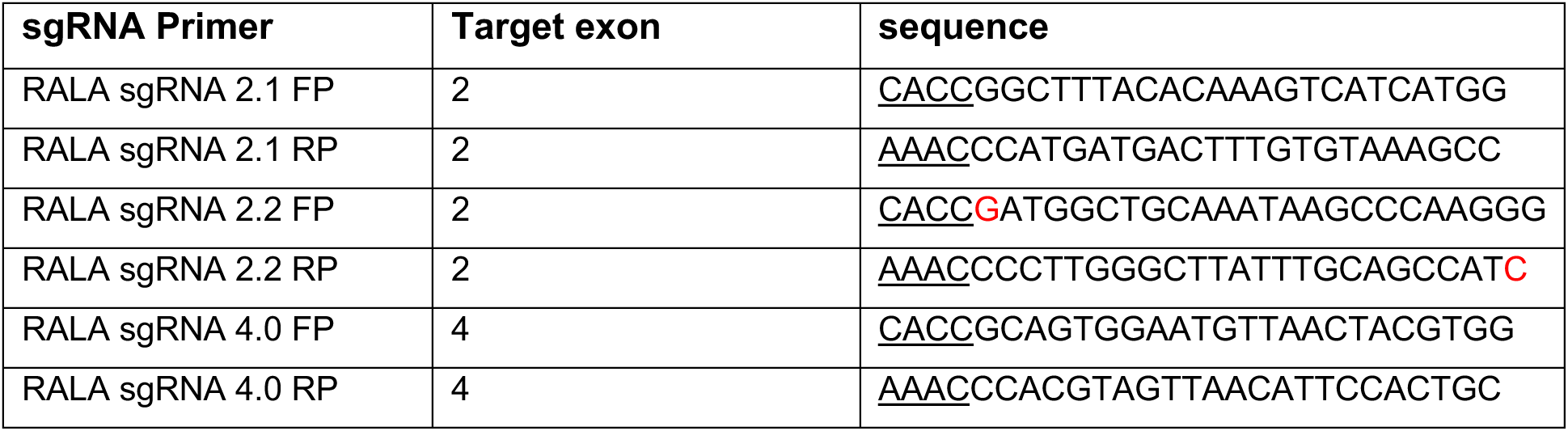
(sgRNA primer design)

To make CRISPR-Cas9 KO, 1 × 10^5^ cells were seeded in 6 cm dish for 24 hours. After 24 hours, cells were transfected with 1ug of empty pSpBBCas9-GFP vector and 1ug of sgRNAs cloned in pSpBBCas9-Puro vector. After 48 hours of transfection, we sorted the cells using GFP as single or double cells in 96 well plates using BD FACSAria cell sorter. Cells were allowed to grow in 10% media till they formed colonies. Colonies were split and cultured in 48 well plates as duplicates. Colonies were screened using western blotting for RALA and Actin. Positive clones were expanded to 6cm dishes, and genotyping confirmed knockout.

### Genotyping of RALA KO cells

For genotyping, cells from a confluent well of 48 well plate were lysed in 200ul of tail lysis buffer and incubated with 2ul of 20mg/ml proteinase K and 2ul of 10mg/ml RNAse A overnight at 55°C (water bath). Proteinase K and RNAse A were denatured by heating at 95°C for 20 min. An equal amount of PCI (Phenol : Chloroform : Isopropanol) was added to this vortex for 1 min and spun at 16000g/4°C/30min. The supernatant was collected in a fresh tube, and an equal amount of Isopropanol was added. The mix was kept at −20°C overnight. Spin at 16000g/4°C/30min and discard supernatant. Wash the DNA pellet with 100ul of ethanol twice at 7500g/4°C/30min and air dry the pellet. Resuspend the DNA in TE buffer or NFW. Use 0.5ug of DNA for genomic PCR in standard conditions. Primers were designed **(as described in table 2)** to bind at the exon-intron boundary and were used to amplify exon 2 and exon 4 (amplicon size: 200bp), and PCR product was resolved on 2% agarose gel.

**Table 2.**
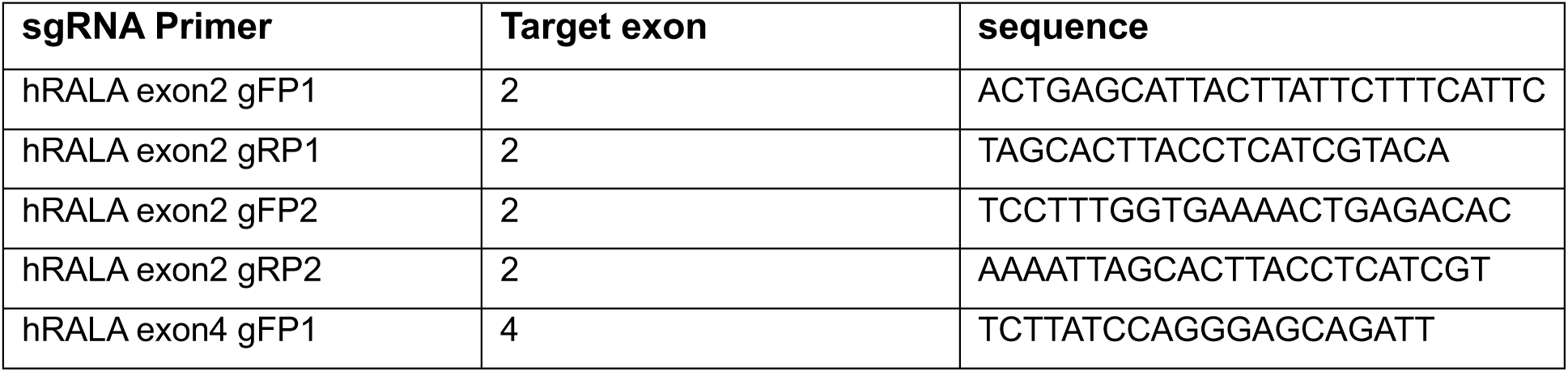
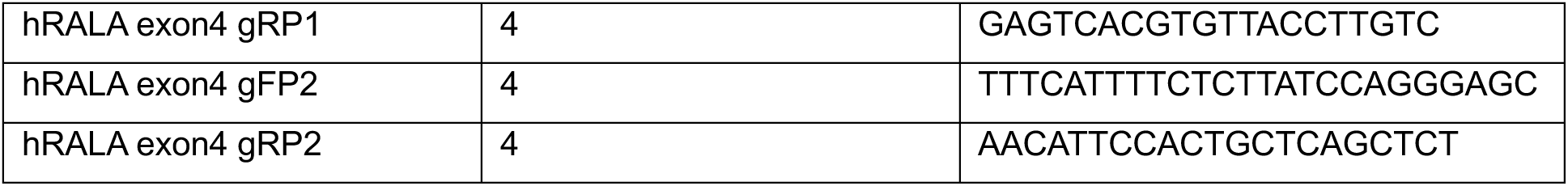
(Genotyping primers)

### Reconstitution of RALA (WT/S194A/S194D) in RALA KO cells

RALA (WT/S194A/S194D) mutants were clones in pBABE puro vector and cloning was confirmed by PCR amplification using reverse primers ending at the mutation site **(as described in Table 3)**.To reconstitute these mutants in RALA KO cells, HEK293T cells were seeded 10 × 10^5^ cells in 10 cm dish. 24 hours after cell seeding, cells were transfected with 2ug of pGagPol, 3ug of pVSVG, 7.5ug of pBABE puro with RALA (WT/S194A/S194D) or empty pBABE GFP and 37.5ul of lipofectamine 2000. 48 hours after transfection, 2ml of fresh media was added to cells. 60 hours after transfection, media (virus titer) was collected and passed through 0.4 micron filter. 1 × 10^5^ RALA KO cells, were seeded in 6 well plate 36 hours after transfection of HEK293T cells. 3ml virus titer, 1ml fresh media and 4ul of polybrene (10mg/ml) were added to the RALA KO cells 24 hours after seeding. Cells were allowed to grow for 48-60 hours (till we see transduction with the pBABE GFP vector). Transduced cells were selected with 1ug/ml puromycin for 48 – 72 hours until all cells in the control plate (transduced with empty pBABE GFP) died. After selection, the media was changed, and cells were allowed to recover for 24 hours. After this, cells were transferred to 24 well plate and western blotting was performed to check the expression of RALA.

**Table 3.**
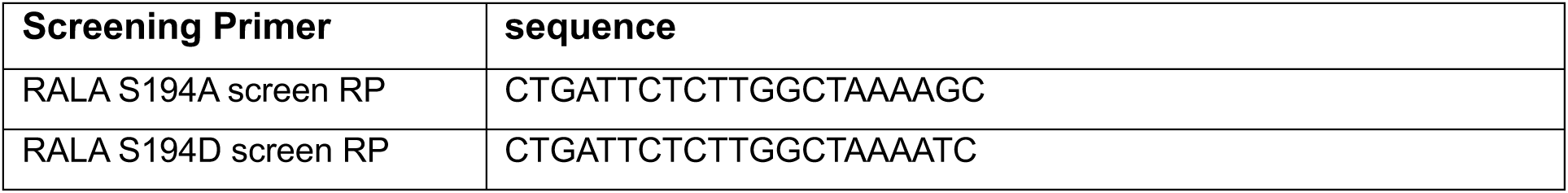
(RALA mutant screening primers)

### Immunofluorescence assay

48 hrs after seeding on glass, cells were fixed with 3.5% paraformaldehyde for 15 minutes. Cells were permeabilized with PBS containing 5% BSA and 0.05% Triton-X-100 for 15 minutes and blocked with 5% BSA for 1 hour at room temperature, followed by incubation with 1:500 mouse anti-RALA (BD Transduction laboratories) antibodies in 5% BSA at 4 degrees for overnight. Cells were finally stained with 1:1000 diluted secondary antibodies (anti-mouse Alexa-488) or 1:500 diluted phalloidin-Alexa 594 for 1 hour at room temperature. All incubations were done in a humidified chamber. Washes were done with 1X PBS at room temperature. Stained and washed coverslips were mounted with Fluoromount-G (Southern Biotech) and imaged using a Zeiss LSM780 multiphoton microscope with a 63x objective.

All image processing was done using FIJI ImageJ software. Initially, the brightness and contrast of all images were adjusted to have a comparable signal intensity. Following this, image thresholding was done to look at the localization RALA in cells.

### siRNA-mediated RALA knockdown in cancer cell lines

Seed 3 × 10^5^ cells per well in 6 well plate for 24 hours. After 24 hours, transfected cells with 50pmol of RALA siRNA **(as described in Table 4)** and 3ul of Lipofectamine RNAimax. After 24 hours, change media and repeat transfection. 48 hours after second transfection, cells were lysed for western blotting to confirm the Kd.

**Table 4.**
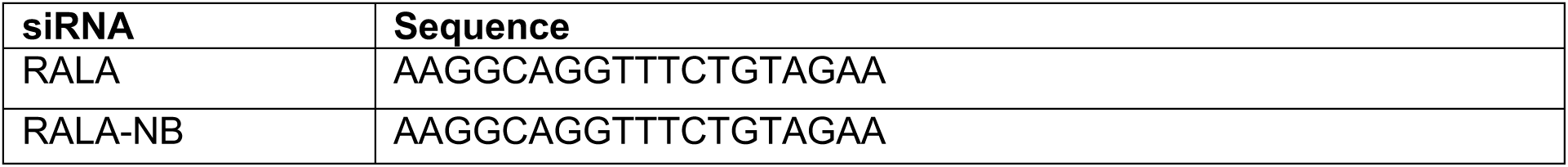
siRNA for RALA Knockdown.

### Mice xenograft

To study the role of RALA and its S194 phosphorylation in tumor formation, we did xenograft assay in an equal number of male and female NOD-SCID mice. 1 million cells were injected subcutaneously in 5-6 week-old mice. Mice were observed over 4-5 weeks, and mice weight and tumor size were measured weekly. At the end of the experiments, mice were euthanized using CO2 and tumors were harvested. 2mm^3^ of the tumor was put in RIPA buffer, and cells were lysed using sonication. Debris was separated by centrifugation at 14000rpm/4°C/15min, and the supernatant was used for protein estimation using the BCA method. Laemmli lysates were prepared, and 20ug of protein was used for western blotting.

### Statistical analysis

All analyses were done using Prism GraphPad analysis software. Statistical data analysis was done using the two-tailed unpaired Student’s T-test, two-tailed paired T-test and, when normalized to respective controls, the two-tailed single sample T-test. Mice tumor data was analyzed using two-way annova (comparing columns within each row).

## SUPPLEMENTARY FIGURE LEGENDS

**Supplementary Figure 1:**

**(A)** DNA sequence alignment of sgRNA clones in pSPCas9(BB)-2A-Puro vector from two colonies (colony1, colony2) each for RALA sgRNA (2.1, 2.2 and 4.0). The sequence for the sgRNA insert (shown in the box) was aligned with the sgRNA sequence used. Overlap is marked by asterix. **(B)** Western blot detection (WB) of AKT, ERK and AMPK in the untreated control (CON), Cas9 control (Cas9), RALA knockout (KO) clones of RAS-independent (MCF7 and SKOV3) and RAS-dependent (T24, UMUC3 and MiaPaCa2) (single selected clone). This was accompanied by the Western blot detection (WB) of active AKT (pAKT – pSer473 AKT), active ERK (pERK – pThr202/Tyr204 ERK1/2) and active AMPK (pAMPK – pThr172 AMPK). **(C)** Genotyping PCR amplicons from untreated control (CON), Cas9 control (Cas9), RALA knockout (KO) clone(s) of RAS-independent (MCF7 and SKOV3) and RAS-dependent (T24, UMUC3 and MiaPaCa2) cells. Exon 2 and 4 were amplified and resolved by agarose gel. Loss of amplicon or change in amplicon size (marked by asterisk) was detected in the gel.

**Supplementary Figure 2:**

**(A)** Uptake of Dextran nano-vesicle with MLN8237+ Rhodamine B (V_MLN+RhB_) in RAS-independent **(MCF7 and SKOV3)** and RAS-dependent **(T24, UMUC3 and MiaPaCa2)** cancer cells were visualized by confocal microscopy in cells treated for 48 hours. The actin cytoskeletal network was stained with phalloidin conjugated to Alexa-488, and the nucleus was counterstained with DAPI.

**Supplementary Figure 3:**

**(A)** DNA sequences for S194A and S194D RALA mutants isolated from positive transformed colonies were compared to known WT RALA sequences (human mRNA NCBI database). The AGT base pair codon for Ser194 is marked in the WT RALA sequence and, when compared, shows distinct mutations in base pair codons for S194A (GCT) and S194D (GAT) mutants. **(B)** WT-RALA, S194A-RALA and S194D-RALA mutants cloned in the pBABE-Puro vector were amplified using RALA primers ending at the mutation site. Primers recognizing WT-RALA (AGT), S194A-RALA (GCT) and S194D-RALA (GAT) were used, and amplicons resolved on agarose gels. **(C)** Western blot detection of RALA S194 phosphorylation (pRALA), total RALA, RALB and Actin in MCF7 control (CON), Cas9 control (Cas9 CON), RALA KO (KO) and RALA KO expressing WT-RALA (WT), S194A-RALA (S194A) and S194D-RALA (S194D) mutants. **(D)** Western blot detection (upper panel) and quantitation (lower panel) of phosphorylation of Threonine 288 residue (pThr288 AURKA), total AURKA from whole cell lysates (WCL) of MCF7 and MIAPaCa2 cells treated for 48hours with DMSO (CON) and empty nano-vesicle scaffold (DEX), 0.02μM Free MLN (MLN) and 0.02μM encapsulated MLN (V_MLN_). The graph represents the ratio of pAURKA/AURKA (normalized to CON as 1) as mean ± SE from three independent experiments. Statistical analysis was done using one sample T-test, and p values, if significant, are represented in the graph (* p < 0.05, ** p<0.01, *** p<0.001).

