## Supplementary Figures for "CRISPR-Cas9 RALA knockout and reconstitution. Detection and role of RALA S194 phosphorylation in RAS-dependent and RAS-independent cancers"

### Supplementary Figure 1

(A)

```
Colony 1  0921_034_002_PLD_SGRALA_2_1_COL2_U6F1_D09.ab1  TCTTGGCTTTATATATCTTGTGGAAAGGACGAAACACCGGCTTTACACAAAGTCATCATG 240
Colony 2  0921_034_001_PLD_SGRALA_2_1_COL1_U6F1_C09.ab1  TCTTGGCTTTATATATCTTGTGGAAAGGACGAAACACCGGCTTTACACAAAGTCATCATG 240
          RALA sgRNA 2.1 sequence                        ----------------------------------CACCGGCTTTACACAAAGTCATCATG 26
                                                                                           **************************

Colony 1  0921_034_002_PLD_SGRALA_2_1_COL2_U6F1_D09.ab1  GGTTTTAGAGCTAGAAATAGCAAGTTAAAATAAGGCTAGTCCGTTATCAACTTGAAAAAG 300
Colony 2  0921_034_001_PLD_SGRALA_2_1_COL1_U6F1_C09.ab1  GGTTTTAGAGCTAGAAATAGCAAGTTAAAATAAGGCTAGTCCGTTATCAACTTGAAAAAG 300
          RALA sgRNA 2.1 sequence                        G----------------------------------------------------------- 27
                                                         *

Colony 1  0921_034_004_PLD_SGRALA_2_2_COL5_U6F1_F09.ab1  GATTTCTTGGCTTTATATATCTTGTGGAAAGGACGAAACACCGATGGCTGCAAATAAGCC 240
Colony 2  0921_034_003_PLD_SGRALA_2_2_COL1_U6F1_E09.ab1  GATTTCTTGGCTTTATATATCTTGTGGAAAGGACGAAACACCGATGGCTGCAAATAAGCC 237
          RALA sgRNA 2.2 sequence                        --------------------------------------CACCGATGGCTGCAAATAAGCC 22
                                                                                               **********************

Colony 1  0921_034_004_PLD_SGRALA_2_2_COL5_U6F1_F09.ab1  CAAGGGGTTTTAGAGCTAGAAATAGCAAGTTAAAATAAGGCTAGTCCGTTATCAACTTGA 300
Colony 2  0921_034_003_PLD_SGRALA_2_2_COL1_U6F1_E09.ab1  CAAGGGGTTTTAGAGCTAGAAATAGCAAGTTAAAATAAGGCTAGTCCGTTATCAACTTGA 297
          RALA sgRNA 2.2 sequence                        CAAGGG------------------------------------------------------ 28
                                                         ******

Colony 1  0921_034_006_PLD_SGRALA_4_COL4_U6F1_H09.ab1  TATTTCGATTTCTTGGCTTTATATATCTTGTGGAAAGGACGAAACACCGCAGTGGAATGT  240
Colony 2  0921_034_005_PLD_SGRALA_4_COL1_U6F1_G09.ab1  TATTTCGATTTCTTGGCTTTATATATCTTGTGGAAAGGACGAAACACCGCAGTGGAATGT  233
          RALA sgRNA 4.0 sequence                      --------------------------------------------CACCGCAGTGGAATGT  16
                                                                                                   ****************

Colony 1  0921_034_006_PLD_SGRALA_4_COL4_U6F1_H09.ab1  TAACTACGTGGGTTTTAGAGCTAGAAATAGCAAGTTAAAATAAGGCTAGTCCGTTATCAA  300
Colony 2  0921_034_005_PLD_SGRALA_4_COL1_U6F1_G09.ab1  TAACTACGTGGGTTTTAGAGCTAGAAATAGCAAGTTAAAATAAGGCTAGTCCGTTATCAA  293
          RALA sgRNA 4.0 sequence                      TAACTACGTGG-------------------------------------------------  27
                                                       ***********
```

(B)

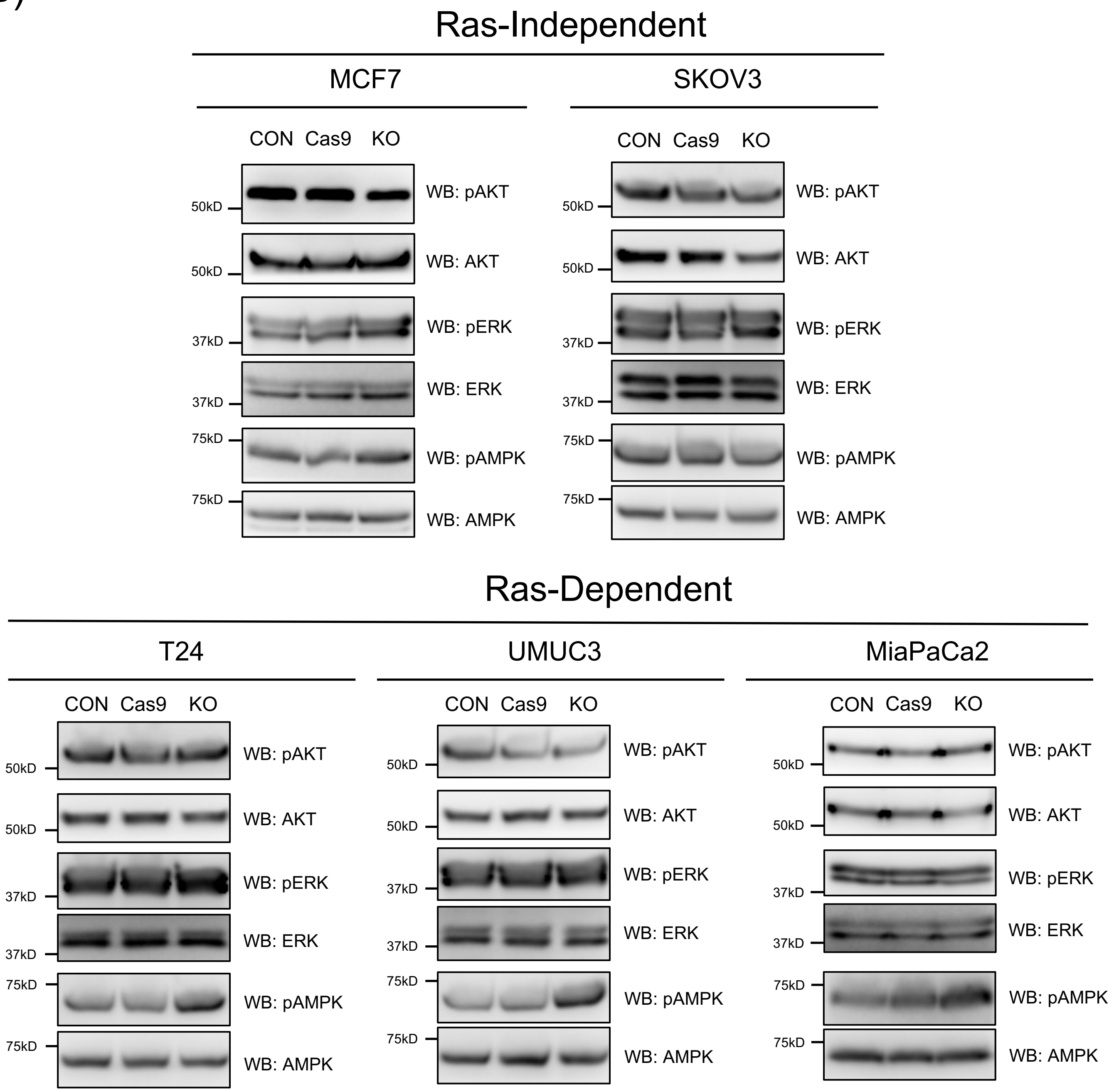

### Supplementary Figure 1

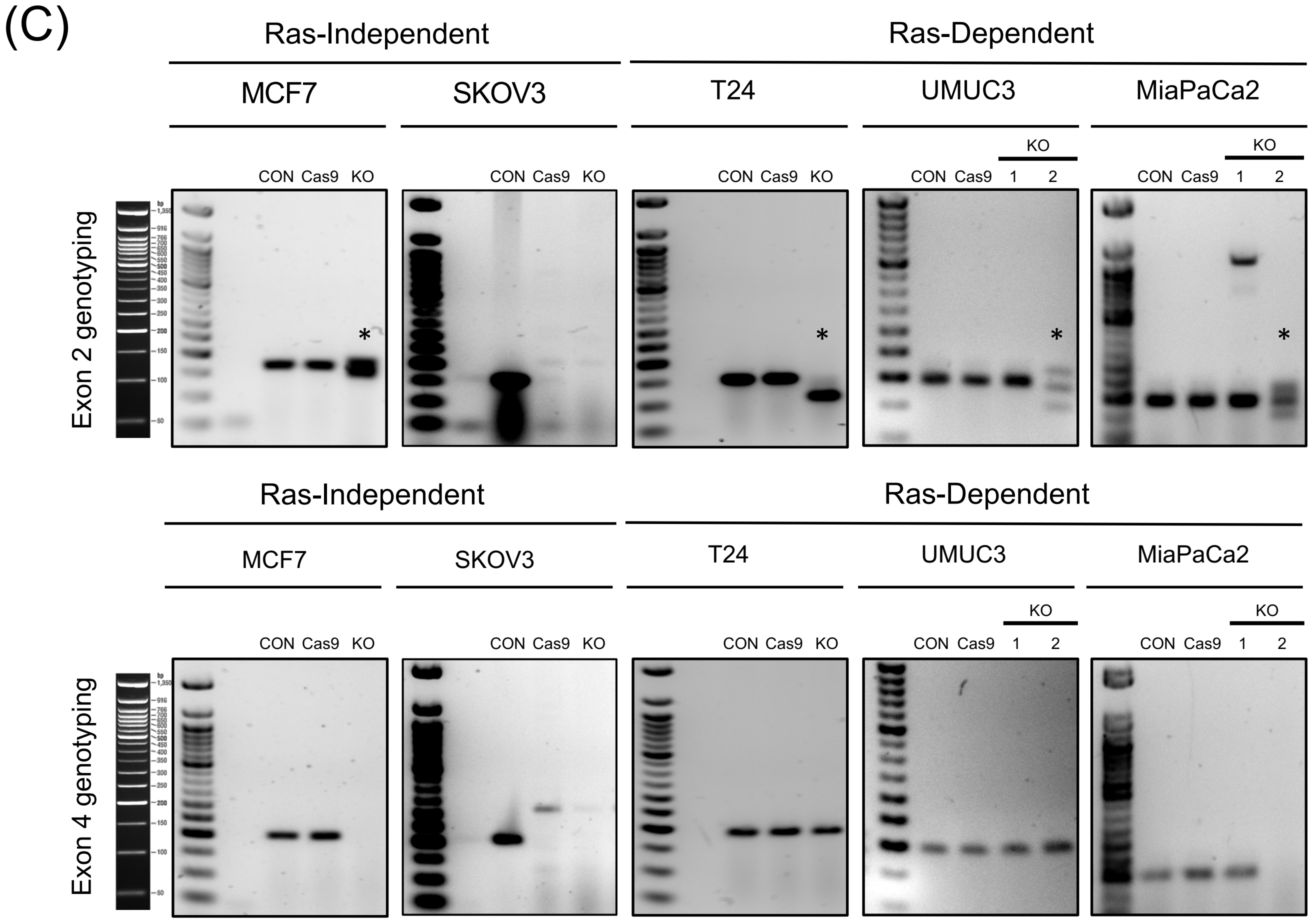

### Supplementary Figure 2

(A)

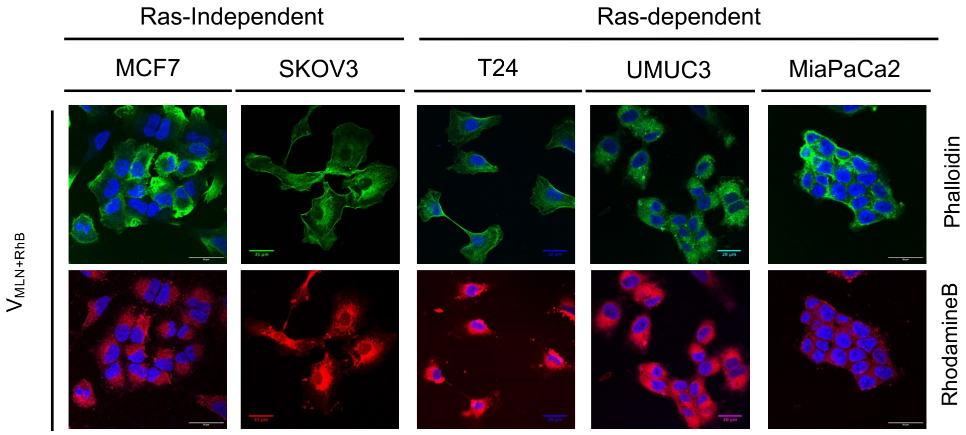

### Supplementary Figure 3:

(A)

RALA S194A cloning (AGT > GCT)

```
RALA WT        545 ACAGCAAAGAAAAGAATGGAAAAAAGAAGAGGAAAAGTTTAGCCAAGAGAATCAGAGAAA 604
S194A Col-2    553 ACAGCAAAGAAAAGAATGGAAAAAAGAAGAGGAAAGCTTTAGCCAAGAGAATCAGAGAAA 612
```

RALA S194D cloning (AGT > GAT)

```
RALA WT        542 AAGACAGCAAAGAAAAGAATGGAAAAAAGAAGAGGAAAAGTTTAGCCAAGAGAATCAGAG 601
S194D Col-2    551 AAGACAGCAAAGAAAAGAATGGAAAAAAGAAGAGGAAAGATTTAGCCAAGAGAATCAGAG 610
```

(B)

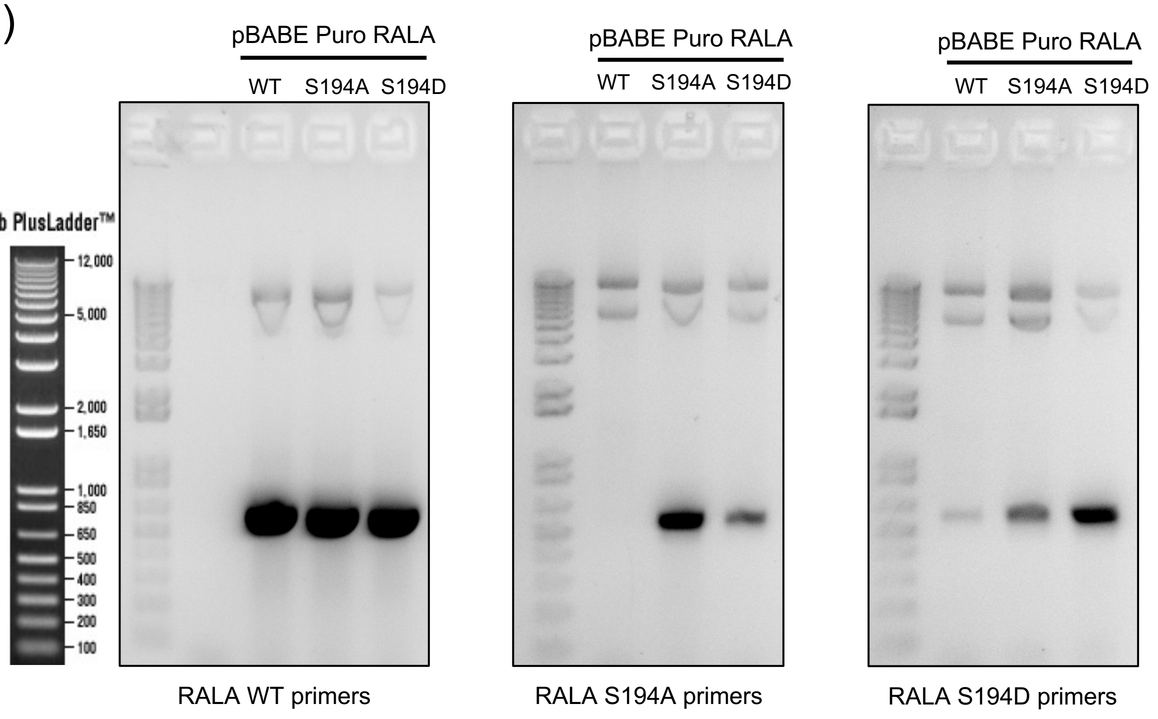


(C)

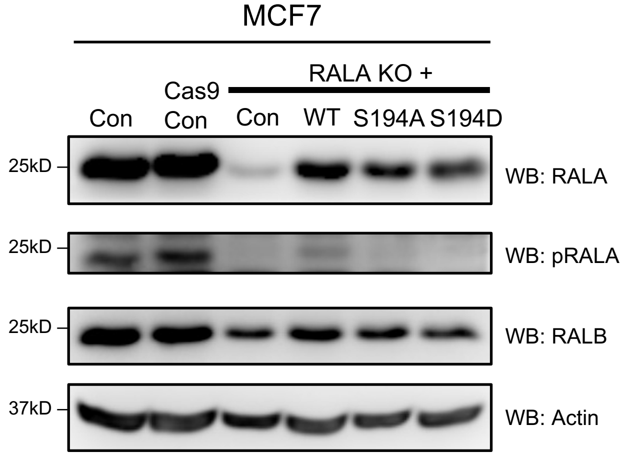

Supplementary Figure 3:

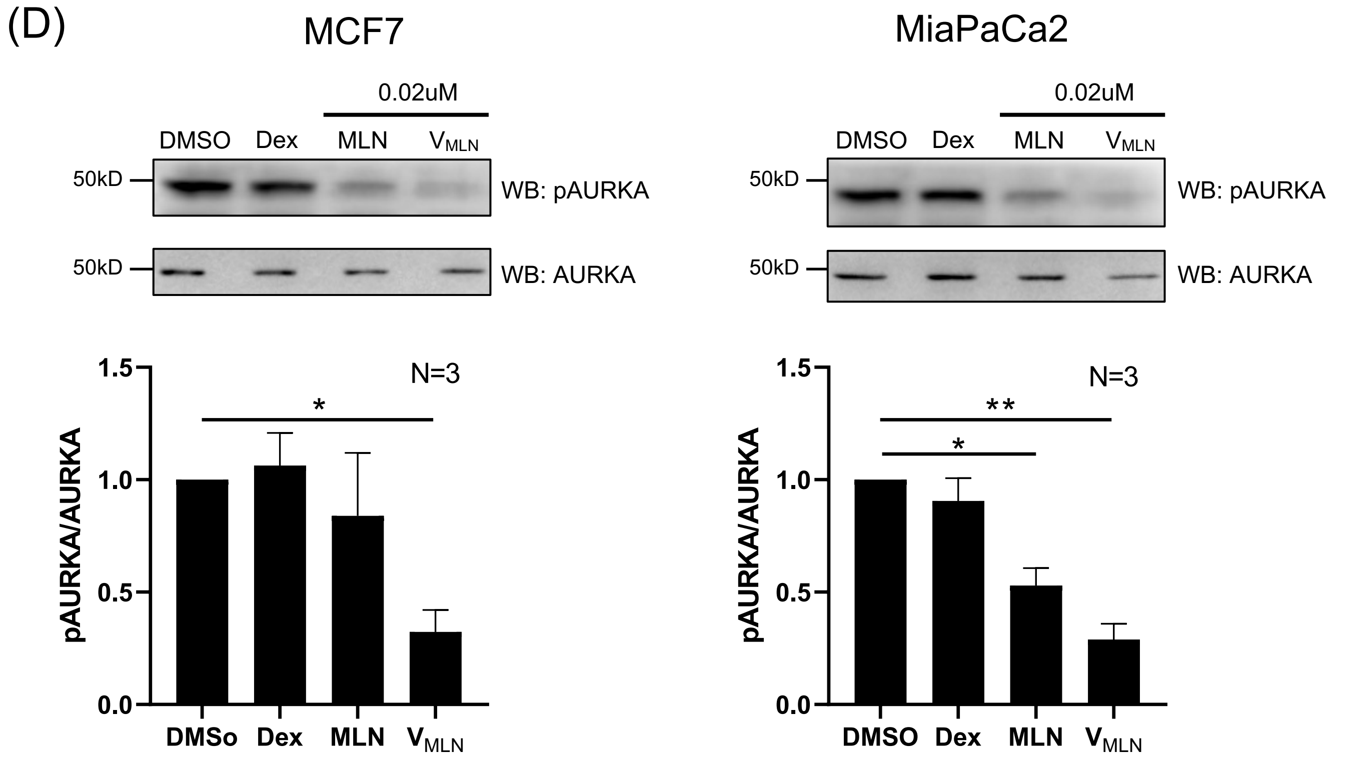
